# Increasing antibiotic susceptibility in *Staphylococcus aureus* in Boston, Massachusetts, 2000-2014: an observational study

**DOI:** 10.1101/125369

**Authors:** Sanjat Kanjilal, Mohamad R. Abdul Sater, Maile Thayer, Georgia Lagoudas, Soohong Kim, Paul C. Blainey, Yonatan H. Grad

## Abstract

**Background:** Methicillin resistant *S. aureus* (MRSA) has been declining over the past decade, but changes in *S. aureus* overall and the implications for trends in antibiotic resistance remain unclear.

**Objective:** To determine whether the decline in rates of infection by MRSA has been accompanied by changes in rates of infection by methicillin susceptible, penicillin resistant *S. aureus* (MSSA) and penicillin susceptible *S. aureus* (PSSA). We test if these dynamics are associated with specific genetic lineages and evaluate gains and losses of resistance at the strain level.

**Methods:** We conducted a 15 year retrospective observational study at two tertiary care institutions in Boston, MA of 31,589 adult inpatients with *S. aureus* infections. Surveillance swabs and duplicate specimens were excluded. We also sequenced a sample of contemporary isolates (n = 180) obtained between January 2016 and July 2016. We determined changes in the annual rates of infection per 1,000 inpatient admissions by *S. aureus* subtype and in the annual mean antibiotic resistance by subtype. We performed phylogenetic analysis to generate a population structure and infer gain and loss of the genetic determinants of resistance.

**Results:** Of the 43,954 *S. aureus* infections over the study period, 21,779 were MRSA, 17,565 MSSA and 4,610 PSSA. After multivariate adjustment, annual rates of infection by *S. aureus* declined from 2003 to 2014 by 2.9% (95% CI, 1.6%-4.3%), attributable to an annual decline in MRSA of 9.1% (95% CI, 6.3%-11.9%) and in MSSA by 2.2% (95% CI, 0.4%-4.0%). PSSA increased over this time period by 4.6% (95% CI, 3.0%-6.3%) annually. Resistance in *S. aureus* decreased from 2000 to 2014 by 0.86 antibiotics (95% CI, 0.81-0.91). By phylogenetic inference, 5/35 MSSA and 2/20 PSSA isolates in the common MRSA lineages ST5/USA100 and ST8/USA300 arose from the loss of genes conferring resistance.

**Conclusions and relevance:** At two large tertiary care centers in Boston, MA, *S. aureus* infections have decreased in rate and have become more susceptible to antibiotics, with a rise in PSSA making penicillin an increasingly viable and important treatment option.

## INTRODUCTION

*Staphylococcus aureus* is one of the most common bacterial pathogens^1,2^. A commensal organism that colonizes the skin and nares of approximately 20% of the population persistently and 60% intermittently^3^, it causes a range of invasive diseases, including skin and soft tissue infection (SSI), bone and joint infection, pneumonia, and bacteremia. Infections are often severe: in 2009, out of 40 million total hospitalizations, an estimated 700,000 were S. aureus-related, yielding a rate of 17.7 per 1,000 hospitalizations^4^.

Much effort has focused on characterizing the emergence and spread of antibiotic resistant *S. aureus* lineages^5^. Resistance to penicillin was documented soon after its introduction and is mediated by a plasmid-borne b-lactamase encoded by the *blaZ* gene^6^. The penicillin resistant phage type 80/81 clone was highly prevalent in the 1950s, before mostly disappearing with the introduction of methicillin in 1960. Subsequently, methicillin resistant *S. aureus* (MRSA) emerged, with resistance conferred by the staphylococcal chromosome cassette *mec* (*SCCmec*), a genetic element that contains the methicillin resistance gene *mecA*. An “archaic” MRSA clone was prevalent until the 1970s and was superseded by the emergence of hospital-associated (HA-MRSA) lineages in the mid to late 1970s, and community-associated (CA-MRSA) lineages in the mid to late 1990s. In the United States the most widesread hospital-associated clonal complex is CC5, which contains the multi-drug resistant MRSA strain known as USA100. Infections acquired in the community are often due to the MRSA strain USA300, which belongs to CC8^7^, though there is evidence for transmission of this lineage in the hospital setting^8^.

The high prevalence of MRSA infections^9,10^ and the increased mortality, cost, and length of hospital stay of individuals infected with MRSA as compared to methicillin susceptible *S. aureus* (MSSA)^11,12^ focused research on describing MRSA epidemiology. In the US and Europe, MRSA incidence peaked in 2005 and has been declining steadily since then^10,13-17^. While the factors driving the decline in MRSA infections are not fully known, infection control efforts in healthcare facilities appear to have reduced the rates of transmission and infection in colonized individuals^16-18^. Evidence from the UK suggests that intrinsic fitness differences between clonal complexes have resulted in a lineage specific decline, possibly due to alterations in selective pressures from changing antimicrobial use^15,16,19-22^.

In contrast with the extensive efforts to characterize MRSA, only a small number of studies have characterized the dynamics of all *S. aureus* subtypes, possibly due to the expectation that prevalence of penicillin susceptible *S. aureus* (PSSA) is extremely low^23^. However, recent reports from diverse sites describe a rising or surprisingly high prevalence of PSSA^24-27^. Whether this trend is associated with the decline in MRSA and how it impacts overall rates of *S. aureus* infection are unclear. Furthermore, recent analyses of large datasets of *S. aureus* genome sequences have provided evidence that resistance is not permanent, but can be acquired and shed^28,29^. However, whether this is generalizable to *S. aureus* lineages present in the United States is unclear.

To determine the overall dynamics of antibiotic resistance in *S. aureus* and evaluate the hypothesis that the decline in rates of MRSA infection has been accompanied by both an absolute and relative increase in PSSA incidence, we analyzed electronic records of *S. aureus* infections in hospitalized patients from 2000 to 2014 at two tertiary care hospitals in Boston, MA. Through whole genome sequencing of contemporary *S. aureus* invasive isolates, we tested the extent to which the trends are associated with specific *S. aureus* lineages and used phylogenomic methods to quantify the gains and losses of penicillin and methicillin resistance.

## METHODS

### Clinical data

Clinical and microbiology information was collected for all inpatients admitted to the Brigham & Women’s Hospital (BWH) and Massachusetts General Hospital (MGH) between January 1, 2000 and December 31, 2014, who were ≥18 years of age and had at least one specimen from any site growing *S. aureus*. Clinical variables included age, gender, Charlson comorbidity index^30^, and the site of infection. Surveillance swabs were excluded. To account for multiple specimens obtained from the same individual, we included the first clinical specimen and excluded specimens with identical antibiograms from the individual collected in the subsequent 2 weeks. We generated two additional datasets using 4 and 6 weeks as the cutoff. Specimens were categorized into four sites of infection: blood, lung (sputum and bronchoscopy specimens), skin and soft tissue (SSI), and other (bone / joint specimens, tissue biopsies, specimens collected from a visceral compartment, urine, and miscellaneous).

### Clinical microbiology

Clinical specimens were analyzed for susceptibility to penicillin (P), methicillin (M), erythromycin (E), clindamycin (C), levofloxacin (L), gentamicin, tetracycline, trimethoprim-sulfamethoxazole (TMP-SMX), rifampin and vancomycin. We define antibiogram-type by the set of antibiotics to which the specimen is resistant. For example, ‘PME’ signifies resistance to penicillin, methicillin, and erythromycin. See Figure S1 of the supplement for the timeline of testing protocols. Susceptibility breakpoints were per Clinical and Laboratory Standards Institute (CLSI) guidelines over the study period; isolates that were ‘intermediate’ by CLSI breakpoints were grouped with resistant isolates for all analyses. We included data for susceptibility to clindamycin after 2010 only since this was the first full year both hospitals performed inducible clindamycin resistance testing on all specimens automatically through the Vitek machine. We categorized a specimen as PSSA if it was susceptible to penicillin and methicillin, MSSA if it was resistant to penicillin but susceptible to methicillin, and MRSA if it was resistant to both penicillin and methicillin.

### Statistical analyses

We analyzed the rate of infection by *S. aureus* per 1,000 inpatient admissions and the mean number of antibiotics to which specimens are resistant. Annual percent changes in counts were adjusted for patient volume. All analyses were stratified by *S. aureus* subtype or antibiogram-type and adjusted for age, gender, Charlson comorbidity index, and site of infection. The analysis with antibiogram-type excluded clindamycin in order to examine trends for the entire study period. Analysis was performed in R version 3.2.2^31^ on data pooled from both facilities. Tests of difference between subtypes or antibiogram-types were 2-sided and comprised of t-tests for continuous variables, chi-squared tests with correction for multiple hypothesis testing for categorical variables, and the Wilcoxon-Mann-Whitney test for the Charlson comorbidity index. Linear and Poisson regression (with patient volume as offset when applicable) were used for multivariable adjustments of rate and count data respectively and 95% confidence intervals were calculated by the profile likelihood method in the R package MASS.

### Prospective specimen collection, sequencing, and analysis

We collected non-duplicate clinical specimens from patients ≥18 years of age who had specimens submitted to the BWH clinical microbiology laboratory between January 1, 2016 and July 22, 2016. The isolates were processed into shotgun sequence libraries on a microfluidic platform using the Illumina Nextera protocol as described^32^ and sequenced on an Illumina platform. A detailed description of isolate preparation is located in the eMethods section of the supplement. We mapped reads to USA300 (GenBank <u>NC_010079.1</u>) by BWA^33^, assembled genomes *de novo* with SPAdes^34^, and annotated them with Prokka^35^. We used Pilon^36^ to call single nucleotide polymorphisms (SNPs). Clonal complex, multilocus sequence types, and *SCCmec* types were assigned using eBURST^37^ and online databases^38,39^. We constructed a maximum likelihood phylogeny from the reference-based SNPs using RAxML^40^ and ST152 (GenBank NZ_LN854556.1) as the outgroup. We inferred by parsimony the number of acquisitions and losses of *SCCmec* within ST5 and ST8 using Mesquite^41^, using N315 (GenBank NC_002745.2) as the reference genome for ST5 and USA300 (GenBank NC_010079.1) as the reference genome for ST8 isolates and with removal of Gubbins-predicted recombination blocks^42^. All genome sequences have been deposited in the National Center for Biotechnology Information Sequence Read Archive (SRA) under SRA accession no. PRJNA380282.

## RESULTS

### Clinical and microbiologic characteristics of S. aureus

Our dataset comprised records of 43,954 *S. aureus* infections, including 21,779 MRSA, 17,565 MSSA, and 4,610 PSSA (Table 1). Patients with MRSA infection were older, had more comorbidities, and more often had lung as the site of infection compared to patients with MSSA or PSSA. Patients with PSSA were older and had more comorbidities compared to those with MSSA but were similar in terms of site of infection. Seventy-eight percent of all specimens belonged to five antibiogram-types (Table S1 of the supplement): two are MRSA (PMEL, PME), two are MSSA (PE, P), and one is PSSA (pan-susceptible). Patients infected by PME *S. aureus* were significantly younger, had fewer comorbidities, and a higher proportion of SSI relative to all other antibiogram-types. Patients with PMEL *S. aureus* infections were significantly older and had a higher proportion of lung infections. Resistance was rare to each of the following: gentamicin, tetracycline, TMP-SMX, rifampin, and vancomycin.

**Table 1:** Demographic and microbiologic characteristics of *S. aureus* subtypes.

|  | All<br>(n = 43,954) | MRSA<br>(n = 21,779) | MSSA<br>(n = 17,565) | PSSA<br>(n = 4,610) | p value* |  |  |
| --- | --- | --- | --- | --- | --- | --- | --- |
|  |  |  |  |  | MRSA vs MSSA | MRSA vs PSSA | MSSA vs PSSA |
| Age (years) | 58.8 (18.3) | 61.4 (18.2) | 55.8 (18.2) | 58.0 (18.1) | <0.0001 | <0.0001 | <0.0001 |
| Female sex | 17,897 (41%) | 9,037 (42%) | 7,023 (40%) | 1,837 (40%) | 0.003 | 0.043 | 0.898 |
| Charlson comorbidity index | 2 (4) | 2 (3) | 1 (3) | 2 (4) | <0.0001 | <0.0001 | 0.001 |
| Race |  |  |  |  |  |  |  |
| Black | 3,463 (8%) | 1,688 (8%) | 1,379 (8%) | 396 (9%) | <0.0001 | 0.0001 | 0.004 |
| Latino | 1,989 (5%) | 831 (4%) | 960 (6%) | 198 (5%) |  |  |  |
| White | 35,083 (84%) | 17,655 (85%) | 13,782 (83%) | 3,646 (83%) |  |  |  |
| Other | 1,220 (3%) | 497 (2%) | 584 (4%) | 139 (3%) |  |  |  |
| Site of infection |  |  |  |  |  |  |  |
| Blood | 6,897 (16%) | 3,187 (15%) | 2,971 (17%) | 739 (16%) | <0.0001 | <0.0001 | 0.018 |
| Lung | 15,110 (34%) | 8,297 (38%) | 5,349 (31%) | 1,464 (32%) |  |  |  |
| SSI | 12,543 (29%) | 5,656 (26%) | 5,518 (31%) | 1,369 (30%) |  |  |  |
| Other | 9,404 (21%) | 4,639 (21%) | 3,727 (21%) | 1,038 (23%) |  |  |  |

Data are mean (SD), n (%), or median (IQR) unless otherwise indicated.
Abbreviations: MRSA, methicillin resistant *S. aureus*; MSSA, methicillin susceptible, penicillin resistant *S. aureus*; PSSA, methicillin and penicillin susceptible *S. aureus*; SSI, skin and soft tissue.
\*Tests of difference are two-sided and comprised of t-test for age; Mann-Whitney test for CCI; chi-squared test for race, gender and site of infection

### Trends in S. aureus infection

After adjusting for covariates including age, site of infection, and comorbidities, rates of infection from *S. aureus* were stable from 2000 to 2003, but subsequently declined annually from 2003 to 2014 by 2.9% (95% CI 1.6%–4.3%; figure 1A and table 2), from 30.6 to 22.0 infections per 1,000 inpatients. This pattern was driven predominantly by MRSA, which declined after 2003 by 9.1% per year (95% CI 6.3%–11.9%) from 20.5 to 5.5 infections per 1,000 inpatients over this time period. In contrast, over the 2003 to 2014 interval PSSA increased 4.6% annually (95% CI 3.0%–6.3%) from 2.1 to 3.6 infections per 1,000 inpatients MSSA declined by 2.2% annually (95% CI 0.4%–4.0%) from 11.7 to 9.2 infections per 1,000 inpatients. There was no change in trends when using 4 and 6 week duplicate exclusion criteria (Figure S2 of the supplement).

**Figure 1:**
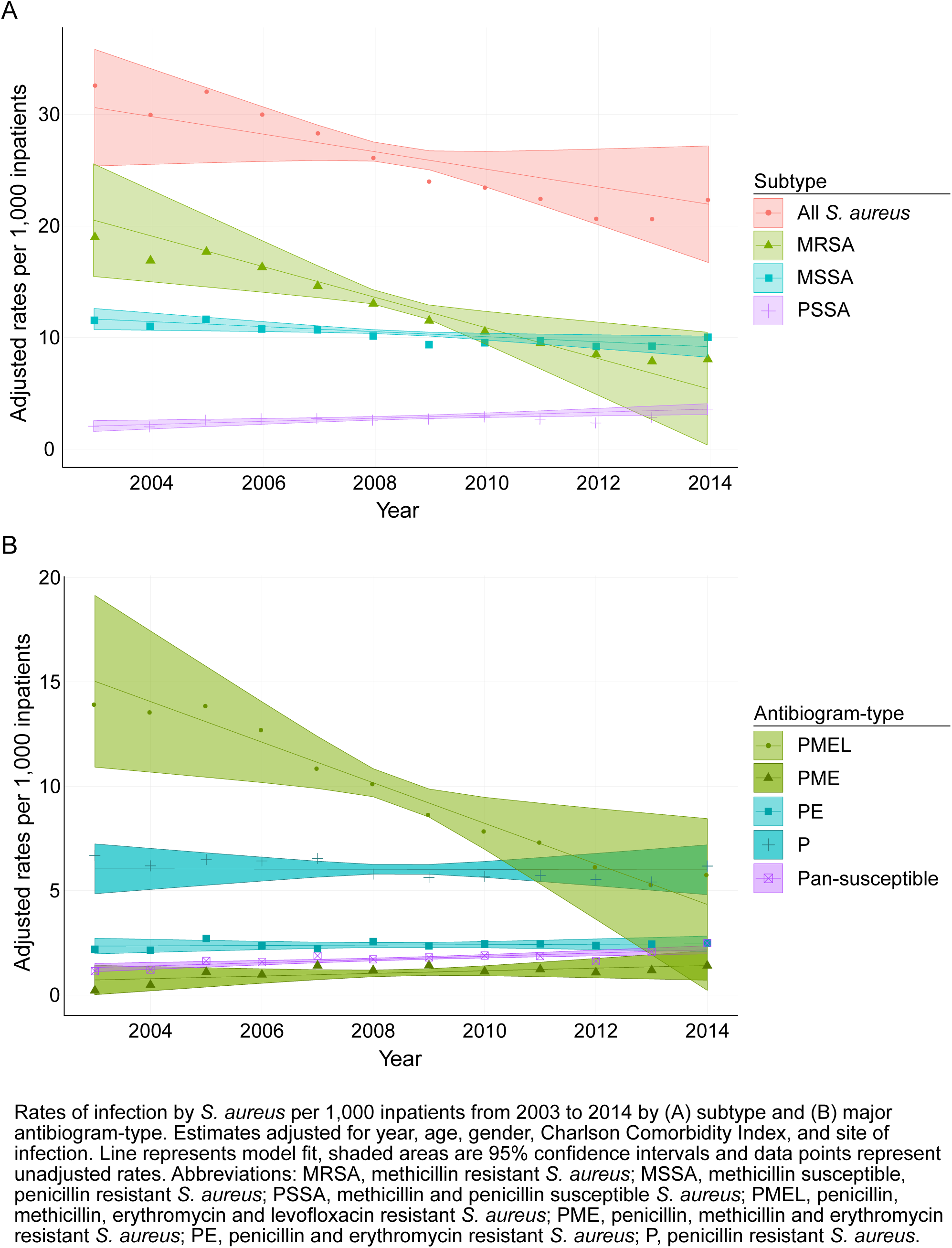
Rates of inpatient *S. aureus* infection by subtype and major antibiogram-type. Rates of infection by *S. aureus* per 1,000 inpatients from 2003 to 2014 by (A) subtype and (B) major antibiogram-type. Estimates adjusted for year, age, gender, Charlson Comorbidity Index, and site of infection. Line represents model fit, shaded areas are 95% confidence intervals and data points represent unadjusted rates. Abbreviations: MRSA, methicillin resistant *S. aureus*; MSSA, methicillin susceptible, penicillin resistant *S. aureus;* PSSA, methicillin and penicillin susceptible *S. aureus;* PMEL, penicillin, methicillin, erythromycin and levofloxacin resistant *S. aureus;* PME, penicillin, methicillin and erythromycin resistant *S. aureus;* PE, penicillin and erythromycin resistant *S. aureus;* P, penicillin resistant *S. aureus*.

**Table 2:** Adjusted rates of inpatient infection of *S. aureus* by subtype and antibiogram type per 1,000 inpatients from 2003 to 2014.

|  | Rate of infection per 1,000 inpatients |  | Annual percent change in counts |  |
| --- | --- | --- | --- | --- |
|  | 2003 (95% CI) | 2014 (95% CI) | Estimate (95% CI) | p value |
| Subtype |  |  |  |  |
| All *S. aureus* | 30.6 (25.4 - 35.8) | 22.0 (16.7 - 27.2) | -2.9 (-4.3 - -1.6) | <0.0001 |
| MRSA | 20.5 (15.5 - 25.6) | 5.5 (0.4 - 10.5) | -9.1 (-11.9 - -6.3) | <0.0001 |
| MSSA | 11.7 (10.7 - 12.6) | 9.2 (8.3 - 10.2) | -2.2 (-4.0 - -0.4) | 0.017 |
| PSSA | 2.1 (1.6 - 2.6) | 3.6 (3.1 - 4.1) | +4.6 (+3.0 - +6.3) | <0.0001 |
| Antibiogram type |  |  |  |  |
| PMEL | 15.0 (10.9 - 19.1) | 4.3 (0.2 - 8.5) | -11.3 (-9.2 - -13.3) | <0.0001 |
| PME | 0.7 (0.0 - 1.4) | 1.4 (0.7 - 2.1) | +6.6 (+2.1 - +11.3) | 0.003 |
| PE | 2.3 (2.0 - 2.7) | 2.4 (2.1 - 2.8) | +0.3 (-1.7 - +2.3) | 0.757 |
| P | 6.0 (4.9 - 7.2) | 6.0 (4.8 - 7.2) | -0.2 (-2.4 - +2.1) | 0.876 |
| Pan-susceptible | 1.3 (1.1 - 1.5) | 2.2 (2.0 - 2.4) | +4.5 (+2.3 - +6.8) | 0.0001 |

Abbreviations: MRSA, methicillin resistant *S. aureus*; MSSA, methicillin susceptible, penicillin resistant *S. aureus*; PSSA, methicillin and penicillin susceptible *S. aureus*; PMEL, penicillin, methicillin, erythromycin and levofloxacin resistant *S. aureus*; PME, penicillin, methicillin and erythromycin resistant *S. aureus*; PE, penicillin and erythromycin resistant *S. aureus*; P, penicillin resistant *S. aureus*.

Rates of infection for the drug-resistant antibiogram-type PMEL declined annually by 11.3% (95% CI 9.2%–13.3%), while rates for the PME antibiogram-type increased annually by 6.6% (95% CI 2.1%–11.3%; figure 1B and table 2). For the two major MSSA antibiogram-types (PE and P), rates did not change significantly over this time period. The pan-susceptible antibiogram-type increased by 4.5% annually (95% CI 2.3%–6.8%). The rates for the PME and pan-susceptible antibiogram-types did not significantly differ. In the analysis including clindamycin for the interval 2010 to 2014, we observed a significant decline in the PMECL antibiogram-type (6.9%, 95% CI 3.7%–10.2%) and a significant increase in the P (3.3%, 95% CI 0.0%–6.6%) and pan-susceptible (10.5%, 95% CI 4.8–16.5%) antibiogram-types (Figure S3 of the supplement).

One explanation for the decline in the MRSA PMEL antibiogram-type is decreasing exposure to antibiotics, particularly fluoroquinolones. We evaluated the rates of resistance to erythromycin, clindamycin and levofloxacin and observed a significant decline for all three drugs in MRSA, a decline for levofloxacin and erythromycin in PSSA, and a decline in erythromycin resistance in MSSA (Figure S4 of the supplement).

### Changes in mean antibiotic resistance

Given the rise in PSSA and the decline in multidrug resistant *S. aureus*, we tested whether overall susceptibility of *S. aureus* changed during the study period. In 2000, a *S. aureus* infection was on average resistant to 3.2 antibiotics. By 2014, this decreased to 2.3 antibiotics in 2014 (p < 0.0001 for trend, figure 2 and table 3). When stratifying by subtype, we found that there was a decline in the mean resistance of MRSA (4.5 antibiotics in 2000 to 3.7 antibiotics in 2014; p < 0.0001 for trend), but there was no change in the average resistance of MSSA and PSSA (p = 0.526 and p = 0.278 for trend, respectively). When including clindamycin for the interval 2010 to 2014, there was an increase in overall resistance but no change in the time trend (Figure S5 of the supplement).

**Figure 2:**
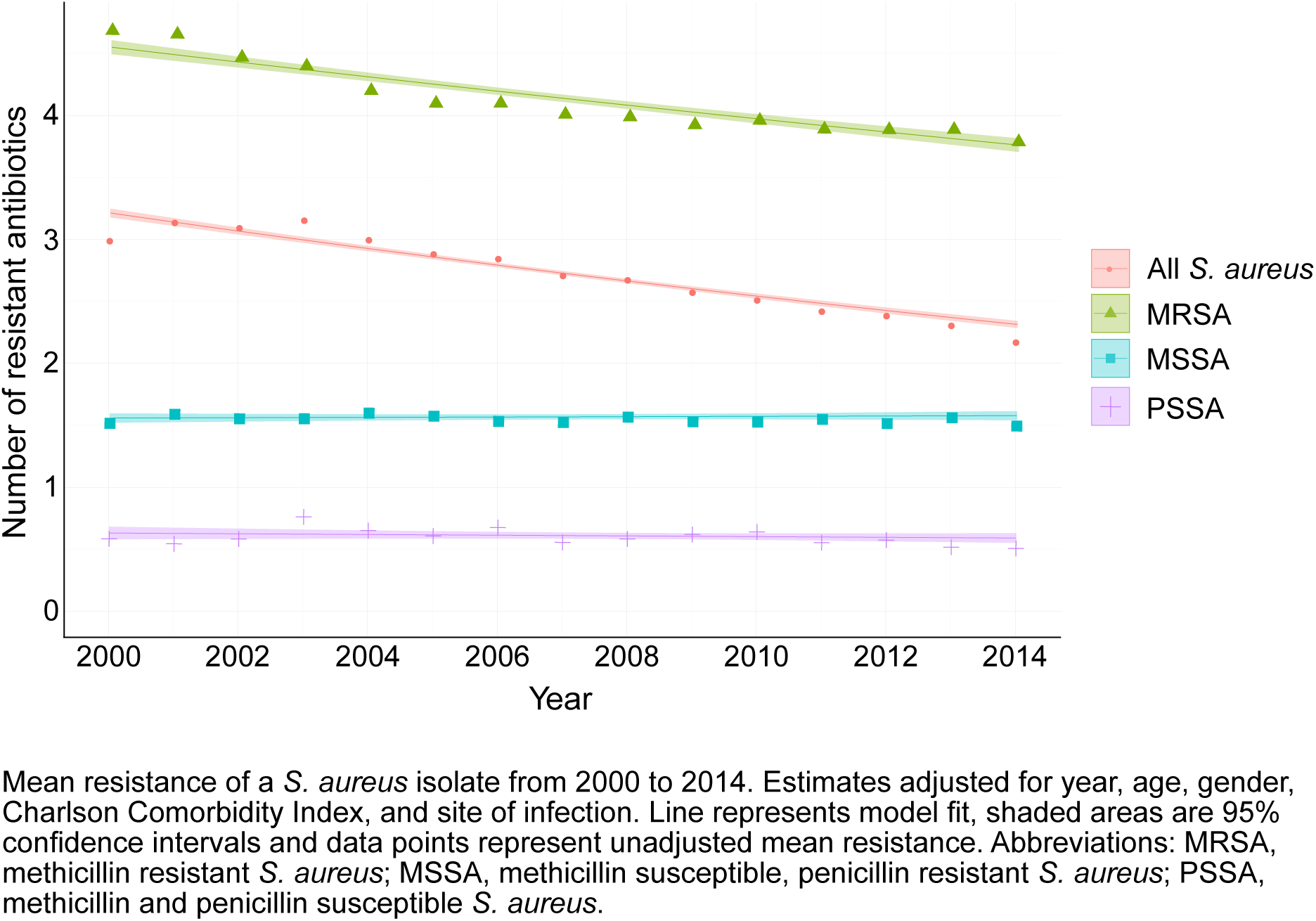
Mean resistance of *S. aureus*. Mean resistance of a *S. aureus* isolate from 2000 to 2014. Estimates adjusted for year, age, gender, Charlson Comorbidity Index, and site of infection. Line represents model fit, shaded areas are 95% confidence intervals and data points represent unadjusted mean resistance. Abbreviations: MRSA, methicillin resistant *S. aureus*; MSSA, methicillin susceptible, penicillin resistant *S. aureus*; PSSA, methicillin and penicillin susceptible *S. aureus*.

**Table 3:** Adjusted mean antibiotic resistance of *S. aureus* by subtype, 2000 to 2014.

|  | Mean resistance (No. of antibiotics) |  | Absolute change (No. of antibiotics) |  |
| --- | --- | --- | --- | --- |
|  | 2000 (95% CI) | 2014 (95% CI) | Estimate (95% CI) | p value |
| Subtype |  |  |  |  |
| All *S. aureus* | 3.2 (3.2 - 3.2) | 2.3 (2.3 - 2.3) | -0.9 (-0.9 - -0.8) | <0.0001 |
| MRSA | 4.5 (4.5 - 4.6) | 3.7 (3.7 - 3.8) | -0.9 (-1.0 - -0.8) | <0.0001 |
| MSSA | 1.5 (1.5 - 1.6) | 1.6 (1.5 - 1.6) | 0.0 (0.0 - +0.1) | 0.526 |
| PSSA | 0.6 (0.6 - 0.7) | 0.6 (0.5 - 0.6) | 0.0 (-0.1 - 0.0) | 0.278 |

Abbreviations: MRSA, methicillin resistant *S. aureus*; MSSA, methicillin susceptible, penicillin resistant *S. aureus*; PSSA, methicillin and penicillin susceptible *S. aureus*.

### Population structure of contemporary S. aureus

Figure 3A illustrates the unrooted phylogeny for a convenience sample of 180 clinical isolates (58 MRSA, 53 MSSA, 69 PSSA), representing 15% of all *S. aureus* isolates identified at the BWH between January 17, 2016 and July 22, 2016. The two most common genetic lineages in this sample were CC5 (n=62) and CC8 (n=43). Other clonal complexes with multiple specimens include CC1 (n=14), CC15 (n=11), CC30 (n=7) and CC45 (n=7). Isolates belonging to each of these CCs clustered together in the phylogeny, with the exception of sequence type 6 (ST6; CC5) and ST72 (CC8) isolates, which were located on separate branches. Thirty-six isolates belonged to minor clonal complexes (each with ≤5 isolates per CC) and 1 isolate had three novel alleles at the MLST loci (Figure S6 of the supplement). MRSA isolates were limited to CC5 (39/62) and CC8 (19/38) (Table S2 of the supplement). Of PMEL isolates, 97% were CC5 and 94% of these were also resistant to clindamycin (PMECL). Of PME isolates, 80% were CC8 and only 20% of these were resistant to clindamycin (Tables S3 and S4 of the supplement). The distribution of gender, comorbidities, race and site of infection differed due to small sample size in the PME group, but otherwise the clinical characteristics of these antibiogram-types were similar between the prospective and retrospective samples (Table S5 of the supplement). MSSA and PSSA isolates and their corresponding antibiogram-types were polyclonal. CC5 and CC8 isolates exhibited a wide range of antibiotic resistance phenotypes (Figure 3B and figure S6 of the supplement). Notably, 23 of 62 isolates (37%) in the hospital-associated lineage CC5 and 24 of 43 (56%) in CC8 were MSSA or PSSA.

**Figure 3:**
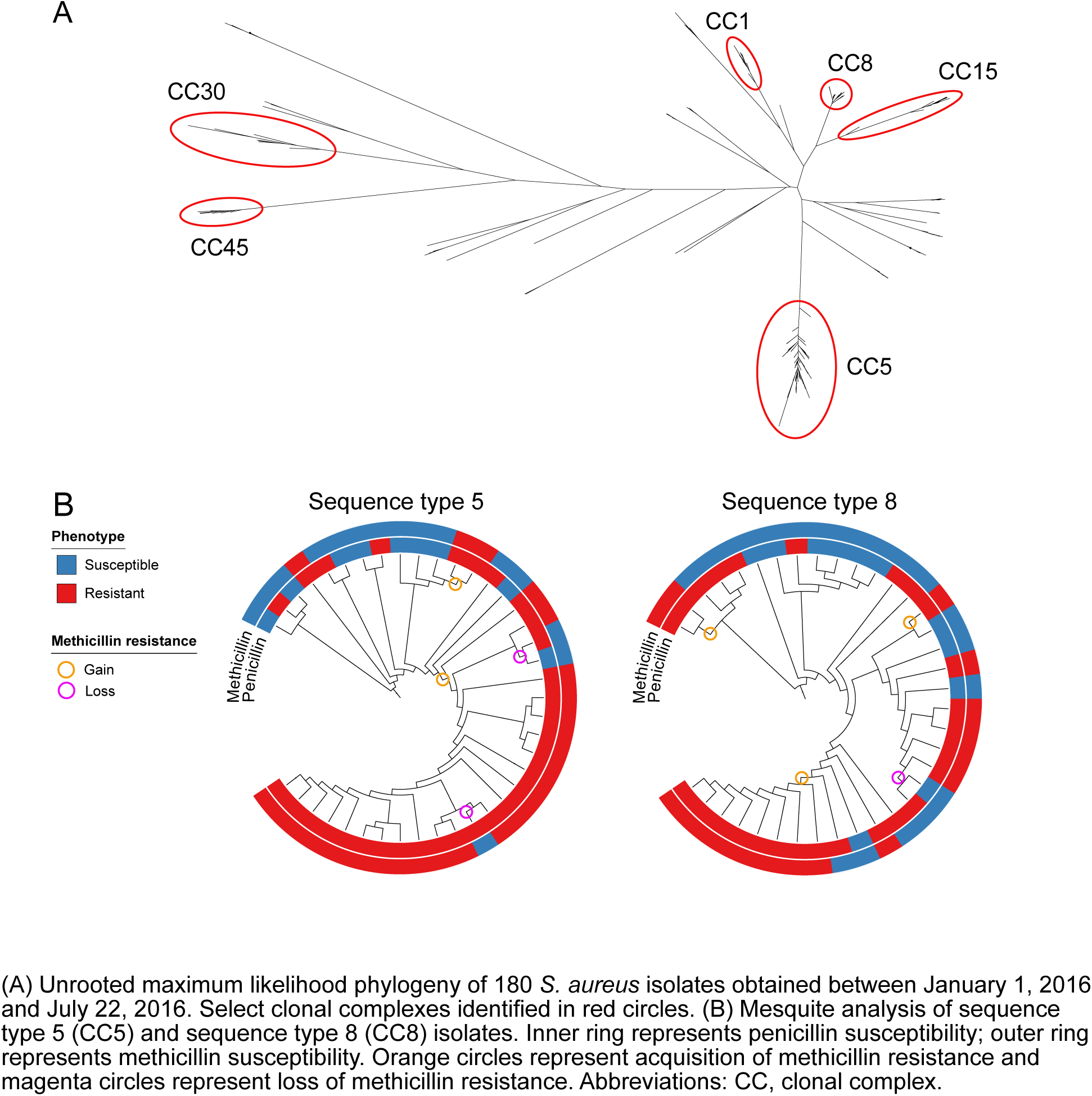
Phylogeny of contemporary *S. aureus* isolates and gain and loss of methicillin resistance at the strain level. (A) Unrooted maximum likelihood phylogeny of 180 *S. aureus* isolates obtained between January 1, 2016 and July 22, 2016. Select clonal complexes identified in red circles. (B) Mesquite analysis of sequence type 5 (CC5) and sequence type 8 (CC8) isolates. Inner ring represents penicillin susceptibility; outer ring represents methicillin susceptibility. Orange circles represent acquisition of methicillin resistance and magenta circles represent loss of methicillin resistance. Abbreviations: CC, clonal complex.

Based on inference from the phylogeny, we identified gain and loss *mecA* (figure 3B), indicating that penicillin and methicillin resistance are dynamic in *S. aureus* populations. In most cases, isolates with loss of *mecA* had complete loss of *SCCmec* (Figure S7 of the supplement). Overall, 3/31 (10%) of PSSA and MSSA in sequence type 8 and 4/24 (17%) in sequence type 5 are inferred to represent isolates that derive from previously MRSA lineages.

## DISCUSSION

Over the past 15 years, the rate of MRSA infection in hospitalized patients at two tertiary care hospitals in Boston, MA declined markedly, the rate of PSSA infection increased, and the overall rate of *S. aureus* infection declined slightly (Figure 1). Combined with the decreased resistance to other antibiotics in MRSA and PSSA, *S. aureus* infections on average have become more antibiotic susceptible over the past 10 years.

The observed decline in the rate of MRSA infection does not reflect a decline in all MRSA strains. As the declining MRSA PMECL antibiogram-type is mostly hospital-associated CC5 and the increasing MRSA PME antibiogram-type is mostly the community-associated CC8, we note that the decline of MRSA appears to be occurring predominantly in strains adapted to the health care setting. The demographic and clinical characteristics of patients with infection from PME isolates support this hypothesis, as they are younger, healthier, and more likely to present with skin and soft tissue infections.

As shown here, studies of MRSA incidence alone fail to capture the dynamics of the overall *S. aureus* population. The observation that penicillin and methicillin resistance are gained and lost at the strain level represents a departure from the dogma that population level resistance is driven by the transmission of lineages with stable resistance phenotypes. Similar findings were noted in a recent study from France that showed *in vivo* loss of methicillin resistance in CC30, a common European lineage that has been on the decline since 2005^29^.

One potential explanation for the loss of resistance across multiple lineages is shifting antibiotic pressures. The decline in use of narrow spectrum beta-lactams such as oxacillin and penicillin G since 2000^43-46^ and in the inpatient use of first generation cephalosporins since 2006^44^ may select against HA-MRSA and in favor of PSSA, but does not on its own explain the rise in CA-MRSA. We also note a decline in levofloxacin resistance in MRSA and PSSA over a time period when use of levofloxacin declined significantly inpatient settings^44^. This suggests that fitness costs associated with fluoroquinolone resistance in *S. aureus* may be a factor in the decline in HA-MRSA and the increase in CA-MRSA and PSSA, which are largely quinolone susceptible. In contrast, use of TMP-SMX increased nationally over the time period of our study^44,47,48^ but TMP-SMX resistance remained <5%. This raises the question whether the fitness cost incurred by TMP-SMX resistance is higher than those incurred by fluoroquinolone resistance.

There are several limitations to our study. First, testing for inducible beta-lactamase production was not routinely performed prior to 2011, raising the possibility that specimens reported as penicillin susceptible from this time period were in fact penicillin resistant. However, such a bias would augment the observed increases in rates of PSSA infection. Second, in the absence of genotyping of historical specimens, the evidence that the decline in MRSA has occurred disproportionately within CC5 relies on the consistency in the demographic and microbiologic characteristics of antibiogram-types between our retrospective and prospective cohorts. However, the over-representation of PMECL isolates in CC5 and PME isolates in CC8 suggests that these antibiogram-types may be rough proxies for lineage. Lastly, the generalizability of our results may be limited as the analysis is from 2 hospitals in the same geographic area. However, as the rate of decline of MRSA in our study is similar to the rate observed in a nationally representative study^13^ and recent reports from geographically diverse institutions note increased rates of PSSA infection, these trends may be widespread. A major strength of this study was the unbiased analysis of the overall *S. aureus* population, whereas most prior studies have examined only the MRSA subpopulation. This has yielded a more complete picture of the clonal dynamics of this highly adaptable pathogen.

The decline in antibiotic resistance in *S. aureus* over the past 10 years runs counter to the prevailing paradigm of an inexorable rise of multidrug resistance among human pathogens. Defining the forces driving the decline will be a critical task to guide efforts to control *S. aureus* infection and antibiotic resistance. Further, as most clinical laboratories do not test for penicillin susceptibility, the increasing incidence of PSSA should prompt evaluation of broader geographic trends along with reevaluation of the cost-effectiveness of penicillin testing and treatment.

## Author contributions

Skanjilal and YHG conceived and designed the study. Electronic medical record data collection and cleaning was done by SKanjilal. Samples were collected by SKanjilal. Samples were processed by GL. Sample preparation / device design and fabrication were done by SKim and PCB. Statistical modeling was performed by SKanjilal and MT. Bioinformatic analysis was done by MAS and YHG. Data interpretation was done by SKanjilal, MAS, and YHG. Drafting of the manuscript was done by SKanjilal, MAS and YHG. All authors critically reviewed and approved the final manuscript.

## Acknowledgements

We thank Marc Lipsitch for advice and discussions and Sarah Fortune for her review of the manuscript. They did not receive compensation for their roles.

